# Reduced intestinal GLP-1^+^ cell numbers are associated with an inflammation-related epithelial metabolic signature

**DOI:** 10.1101/2025.03.05.641577

**Authors:** Elisabeth Urbauer, Doriane Aguanno, Katharina Kuellmer, Amira Metwaly, Nadine Waldschmitt, Mohamed Ahmed, Sevana Khaloian, Gabriele Hörmannsperger, Julien Planchais, Tobias Fromme, R Balfour Sartor, Harry Sokol, Dirk Haller, Eva Rath

## Abstract

**Background & Aims:** Enteroendocrine cells (EECs) are known for their role in digestion and metabolism, yet their role in intestinal inflammation remains unclear. In inflammatory bowel diseases (IBD), a contribution of EECs to pathogenesis is indicated by autoantibodies affecting EEC function and general disease symptoms like insulin resistance and altered intestinal motility. Particularly, the L cell-derived hormone glucagon-like peptide 1 (GLP-1), suggested to orchestrate metabolic-inflammatory responses may influence inflammatory pathways in the intestine.

**Methods:** We quantified numbers of GLP-1^+^ cells in 4 different mouse models of intestinal inflammation and performed transcriptional analyses of colonic epithelial cells from inflamed interleukin (IL)10-deficient mice. Using a publicly available single-cell RNA sequencing dataset including mucosal biopsies from Crohn’s disease (CD) patients, we confirmed findings from the murine models. A model of mitochondrial dysfunction (ClpP^ΔIEC^ mice) as well as murine and human intestinal organoids were used to study molecular mechanisms.

**Results:** Numbers of GLP-1 expressing cells are consistently reduced at the site of active disease in mouse models and CD patients. Despite this reduction, L cells from inflamed IL-10-deficient mice remained functional regarding GLP-1 secretion. Transcriptional analyses of intestinal epithelial cells indicate altered differentiation correlating with an inflammatory metabolic fingerprint. Reduced GLP-1^+^ cells in ClpP^ΔIEC^ mice and inhibition of respiration in organoid cultures supports a causative role for metabolism in steering differentiation.

**Conclusion:** Reduction of GLP-1^+^ cells represents a general feature of ileal and colonic inflammation in mice and human. Given the numerous properties of GLP-1, this reduction likely affects inflammatory processes in the mucosa and disease-related symptoms on multiple levels, and therefore, should be considered a therapeutic target in IBD.

**Data Transparency:** All data generated or analyzed during this study are included in this published article. Additional datasets, including raw data, are available from the corresponding author upon reasonable request.

**Synopsis:** This study examines GLP-1^+^ cells in intestinal inflammation, showing consistent reductions in inflamed areas. Findings from mouse models and human data reveal an inflammatory metabolic profile linked to altered epithelial differentiation. GLP-1, involved in endocrine-immune crosstalk, may impact mucosal inflammation and symptoms, making it a therapeutic target.

**Graphical Abstract:** 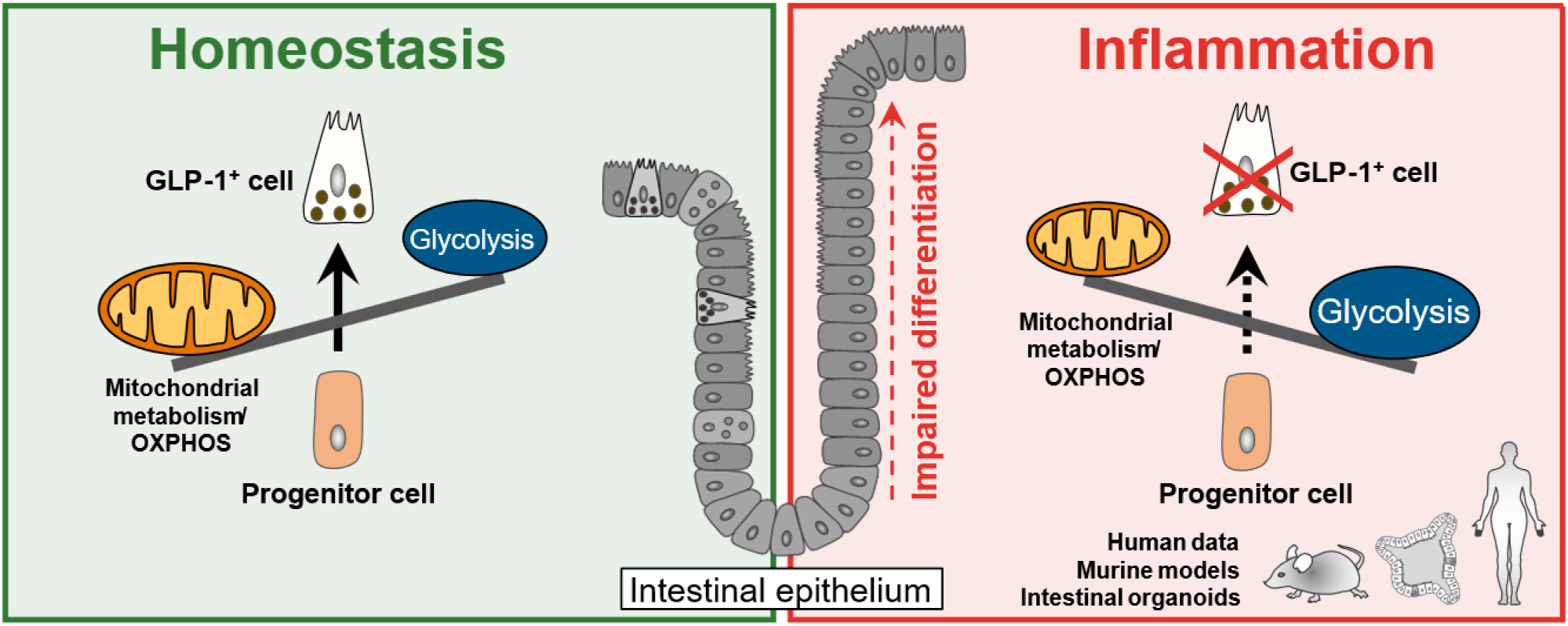

## 1 Introduction

Enteroendocrine cells (EECs) are renowned for regulating gastrointestinal motility, secretion of digestive fluids, and inducing insulin release *via* peptide hormones. In particular, glucagon-like peptide 1 (GLP-1) has gained enormous attention due to its insulinotropic action and relevance in the treatment of type 2 diabetes and obesity (1). EECs constitute approximately 1% of intestinal epithelial cells (IECs) and are scattered throughout the intestine (2). The intestinal epithelium itself is a key player in maintaining metabolic and immune homeostasis. Ensuring efficient nutrient absorption and at the same time forming a barrier for host defense (3), IECs constitute the bodýs most important and dynamic interface to the external environment (4). Stem cells localized at the crypt base expressing leucine-rich repeat-containing G-protein coupled receptor 5 (Lgr5) constantly divide and renew the IEC layer within 3-5 days (5), giving rise to different subtypes of IECs. These subtypes, comprising enterocytes, Paneth cells, goblet cells, tuft cells, M cells, and EECs (6), fulfill specialized functions, with absorptive enterocytes and secretory (mucin-producing) goblet and (antimicrobial peptides-producing) Paneth cells accounting for the largest proportion of IECs.

The regulation of stem cell renewal, proliferation and differentiation is of particular importance under pathological conditions, such as wound healing, tissue regeneration and acute and chronic inflammation as seen in inflammatory bowel diseases (IBD). IBD comprises ulcerative colitis (UC) and Crohn’s disease (CD), which are characterized by relapsing inflammation of the colon (UC) or different areas of the gastrointestinal tract but predominantly affecting the terminal ileum (CD). The current paradigm for IBD pathogenesis is a dysregulated interaction between the intestinal microbiota and the mucosal immune system. Consequently, failure of IECs as mediators between these two major factors influencing intestinal inflammation (3) may critically contribute to IBD pathogenesis (7).

Alterations in IEC subpopulations and aberrances of the intestinal stem cell niche have been described in the context of intestinal inflammation. The reduction of secretory goblet cells and Paneth cells (7–9) are the best-studied phenomena in this context. However, upon parasite infection increases in tuft cells, CCK-positive cells, and serotonin (5-HT)-producing enterochromaffin cells are also observed (10–12). Yet, GLP-1-expressing L cells as additional members of the secretory linage of IECs have not been thoroughly characterized under inflammatory conditions.

Loss of goblet cells and Paneth cells are early events in intestinal inflammation (9, 13) and have been associated with impaired metabolic fitness of IECs (9, 14). In general, the cellular metabolism plays a pivotal role in regulating intestinal epithelial cell proliferation and differentiation, and mitochondrial function actively determines cellular phenotypes and lineage commitment (9, 15–17). Consistently, there is extensive evidence for altered cellular metabolism, mitochondrial dysfunction, and activation of mitochondrial stress signaling in IBD (15, 18, 19). Moreover, genetic risk loci associated with IBD have been mapped to mitochondrial function-associated genes (15, 20–23) and mitochondrial dysfunction-associated aberrances of the intestinal stem cell niche predict disease recurrence in CD patients (9), suggesting alterations in epithelial cell oxidative metabolism not only as a consequence but causal factor in the development of chronic intestinal inflammation.

Even though EECs have been identified as source of the disease-relevant cytokine IL-17C in IBD (24), and autoantibodies against the enteroendocrine protein UbE4A are associated with disease behavior (25), studies investigating the role of EECs in IBD pathogenesis are scarce. Yet, particularly GLP-1 producing cells might be of interest in the context of inflammation as these cells actively sense the microbiota and bacterial products, and in addition to its role in glucose metabolism, GLP-1 exerts potent anti-inflammatory effects (26, 27). Highlighting the crosstalk between the immune and endocrine system, the GLP-1 receptor is expressed on multiple immune cell populations, including intestinal intraepithelial lymphocytes (28, 29). In addition, general disease symptoms including reduced appetite, anorexia and altered intestinal motility often accompanying intestinal inflammation, might be linked to GLP-1 signaling (2, 30, 31).

As the total number of studies investigating GLP-1^+^ cell numbers and function in the context of intestinal inflammation is limited and results are conflicting (26, 27, 32–34), we quantified numbers of GLP-1 expressing cells in 4 different mouse models of ileal and colonic inflammation (*Tnf^ΔARE^*mice (ileitis), IL-10^-/-^ mice and adoptive T cell-transfer model (colitis) as well as *Citrobacter rodentium*-infection (colitis)). Consistent across all models, a reduction of GLP-1^+^ cells confined to the site of active inflammation is evident, which occurs early in the course of inflammation. The remaining GLP- 1-expressing L cells in IL-10^-/-^ mice are functional in terms of hormone secretion. Importantly, analysis of a recently published data set from single-cell RNA sequencing of mucosal biopsies from CD patients (35), confirmed the human relevance of these findings. In parallel to EEC loss, colonic IECs of inflamed IL-10^-/-^ mice display a metabolic fingerprint that has been associated with reduced differentiation into other cell types of the secretory lineage (goblet cells, Paneth cells). Remarkably, compromised mitochondrial function in ClpP^ΔIEC^ mice, as well as inhibition of mitochondrial respiration using oligomycin in murine and human organoid cultures was sufficient to reduce EEC marker gene expression, indicating a causative role for metabolism in steering differentiation into this IEC subtype.

## 2 Results and discussion

### 2.1 GLP-1 positive cells are consistently reduced under inflammatory conditions

To comprehensively characterize the impact of intestinal inflammation on the number of GLP-1- expressing L cells in the epithelium, ileal and colonic tissue sections from different mouse models of intestinal inflammation were immunohistochemically stained (**Figure 1A**). Cells positive for GLP-1 were quantified as positive cells over the total epithelial layer (mm^2^), to minimize confounders due to architectural changes of the intestinal tissue caused by inflammatory processes such as immune cell infiltration. At least 3 non-consecutive, well-oriented tissue sections per mouse were used for quantification. Group sizes were ≥ 5 in all cases. In accordance with 3R principles, tissue samples analyzed were obtained from our tissue biobank and originated from previous studies (9, 36–38). Mouse models investigated comprise commonly used models of IBD, (I) the *Tnf^ΔARE^* mouse model of ileitis and (II) IL-10 deficient mice as model of colitis; (III) an adoptive T cell transfer model of colitis, Rag2^-/-^ mice reconstituted with CD4^+^ CD25^-^ T cells; and (IV) an infection model using *Citrobacter rodentium*. In *Tnf^ΔARE^* mice, CD-like tissue inflammation develops gradually and microbiota-dependent due to loss of translational control of the pro-inflammatory cytokine tumor necrosis factor (Tnf) caused by a deletion of AU-rich (adenosin-uracil) elements (ARE) in the *Tnf* gene/ mRNA, resulting in Tnf overexpression (39). Similarly, mice lacking the anti-inflammatory cytokine interleukin 10 (IL-10^-/-^ mice) spontaneously develop colitis after weaning whereby onset and severity of inflammation are dependent on the microbiota and the genetic background of mice (40), with the 129/SvEv mouse line being more susceptible than C57BL/6 mice. In the following, “IL-10^-/-^ mice” refers to inflamed 129/SvEv IL-10 deficient mice unless otherwise stated. The T cell transfer model of colitis is the most widely used model of IBD to dissect the contribution of T cells to immunopathology in chronic colitis (41). Notably, the expansion of enterochromaffin cells, a cell type closely related to EECs, is dependent on T cell presence during helminth infection (10), indicating a direct effect of immune cells on epithelial cell composition. In contrast to the models of chronic inflammation, infection with *C. rodentium* causes transient colonic inflammation and hyperplasia and serves as a model to study host- pathogen interaction in the context of IBD and to study molecular mechanisms of human gastrointestinal infections with pathogenic *Escherichia coli* strains (42).

**Figure 1:**
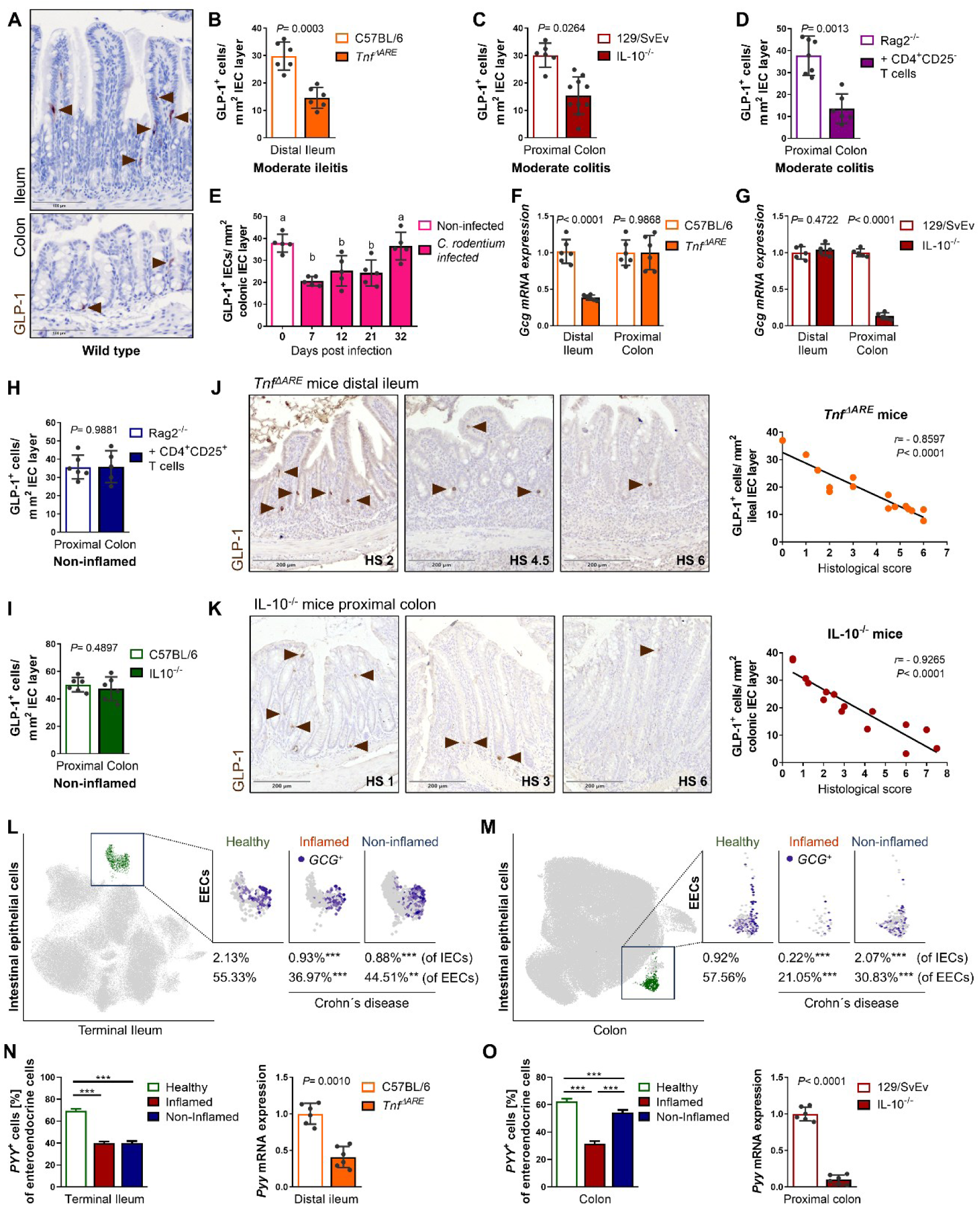
GLP-1^+^ cells are reduced in the murine and human intestinal mucosa under inflammatory conditions. (**A**) Representative pictures of ileal and colonic tissue sections from wild type mice immunohistochemically (IHC) stained for GLP-1. Arrows indicate cells positive for GLP1. (**B-D**) Quantification of GLP-1-positive cells (**B**) in ileal tissue sections of moderately inflamed (histological score (HS) 2-4) *Tnf^ΔARE^*mice, (**C**) in colonic tissue sections of moderately inflamed IL-10^-/-^ mice, (**D**) in colonic tissue sections of moderately inflamed Rag2^-/-^ mice adoptively transferred with colitogenic CD4^+^ CD25^-^ T cells, and respective controls. (**E**) Quantification of GLP-1-positive cells in colonic tissue sections of mice infected with *Citrobacter (C.) rodentium* prior to and at different time points post infection. (**F, G**) qRT-PCR analysis for *Gcg* (encoding GLP-1) of ileal and colonic tissue from (**F**) *Tnf^ΔARE^* mice and (**G**) from IL-10^-/-^ mice including respective controls. Expression levels are normalized to controls and tissue of origin. (**H, I**) Quantification of GLP-1-positive cells in colonic tissue sections (**H**) of non-inflamed Rag2^-/-^ mice adoptively transferred with regulatory CD4^+^ CD25^+^ T cells and (**I**) of non-inflamed IL-10^-/-^ mice on C57BL/6 background and respective controls. (**J, K**) Representative GLP-1 IHC stainings for the HS indicated (left) and correlation analysis (Pearson) of the number of GLP-1-positive cells and HS for (**J**) ileal tissue from *Tnf^ΔARE^* mice and (**K**) colonic tissue from IL-10^-/-^ mice. (**L**, **M**) Uniform manifold approximation and projection (UMAP) visualization of scRNA-sequencing of intestinal epithelial cells from the terminal ileum (**L**) and colon (**M**) from Crohn’s disease patients (35). The subgroup of enteroendocrine cells is highlighted in green (left) and *GCG*-expressing enteroendocrine cells are shown in purple in healthy, inflamed and non-inflamed conditions (right). The sample size for healthy/ inflamed/ non-inflamed is 11/ 14/ 45 for (**L**) and 32/ 5/ 18 for (**M**), respectively. Chi-square test and Fisheŕs exact test, asterisks indicate differences compared to healthy controls, \**P*<0.05, \*\**P*<0.01, \*\*\**P*<0.001. (**N, O**) Left: Barplots showing the proportion of EECs expressing *PYY* in the terminal ileum **(N**) and colon (**O**) in the same cohort of CD patients and controls. Bars represent mean+SD, One-way ANOVA followed by Tukey test, asterisks indicate significant differences \**P*<0.05, \*\**P*<0.01, \*\*\**P*<0.001. Right: qRT-PCR analysis for *PYY* of ileal tissue from *Tnf^ΔARE^* mice (**N**) and of colonic tissue from IL-10^-/-^ mice (**O**) and respective controls. (**B-I, N, O**) Bars represent mean+SD, n≥5 for all groups. Unpaired t tests except (**E**) one-way ANOVA followed by Tukey test. *P* values and r are indicated; a is significantly (*P* < 0.05) different from b.

In a first approach, we compared moderately inflamed mice (histopathological score (HS) 2-4) of the different models to wild type (Wt) littermates or non-reconstituted/ non-infected mice, respectively (**Figure 1B-E**). Consistently throughout all models investigated, numbers of GLP-1^+^ cells were reduced under inflammatory conditions (**Figure 1B-E**). These changes were observed at the site of active disease, namely the ileum in *Tnf^ΔARE^*mice (**Figure 1B**) and the colon in IL-10^-/-^ mice, Rag2^-/-^ mice adoptively transferred with colitogenic CD4^+^ CD25^-^ T cells, and C57BL/6 mice infected with *C. rodentium* (**Figure 1C-E**). Of note, during infection with *C. rodentium*, loss of GLP-1^+^ cells was transient and paralleled tissue aberration in the progressive and regressive phases of infection, with protein expression returning to basal numbers after bacterial clearance (day 32) (**Figure 1E**). Reflecting cell numbers, we observed diminished expression levels of *Gcg* (*glucagon*, translated *i.a.* into GLP-1 in the intestine) at the site of active disease in *Tnf^ΔARE^* mice and IL-10^-/-^ mice but not unaffected tissue regions (proximal colon and distal ileum, respectively) (**Figure 1F,G**), indicating a local and acute effect of inflammatory processes on intestinal epithelial cell composition. Loss of IL-10 receptor has been shown to disrupt IEC proliferation and skew differentiation towards the goblet cell fate, implicating a role for IL-10 signaling in IEC lineage decisions (43). Yet, corroborating an inflammation-mediated over a genetic effect, we did not find any reduction in GLP-1^+^ (**Figure 1H,I**) in non-inflamed IL-10^-/-^ mice on C57BL/6 background or Rag2^-/-^ mice reconstituted with regulatory CD4^+^ CD25^+^ T cells, thus not developing colitis. Notably, the reconstitution of Rag2^-/-^ mice with regulatory T cells did not exert any effect on GLP-1^+^ cell numbers (**Figure 1H**), suggesting lack of T cell-mediated regulation under homeostatic conditions.

For characterizing the effect of inflammation on EECs numbers in detail, we took advantage of the heterogeneity in the grade of CD-like inflammation in *Tnf^ΔARE^* mice and of colitis in IL-10^-/-^ mice. In independent, additional cohorts, we quantified numbers of GLP-1^+^ across a broad range of histopathological scores. Numbers of cells positive for GLP-1 inversely correlated with the grade of inflammation in the ileum of *Tnf^ΔARE^* mice (**Figure 1J**) and in the colon of IL-10^-/-^ mice (**Figure 1K**). Comparing tissues from non-inflamed (HS <2) *Tnf^ΔARE^* mice and Wt C57BL/6 littermates as well as IL-10^-/-^ mice and Wt 129/SvEv controls, we observed no differences in the total numbers of GLP-1^+^ (**Figure 1B,C,J,K**). Thus, loss of EECs parallels the progression towards an inflammatory tissue environment.

To confirm the human relevance of our findings, we used a recently published single-cell RNA sequencing data set from mucosal biopsies comprising healthy controls, as well as inflamed and non-inflamed tissues from CD patients (35). A total of 154,136 epithelial cells derived from terminal ileum (**Figure 1L**) and 97,788 colonic epithelial cells (**Figure 1M**), respectively, were clustered into subgroups and EECs were analyzed for *GCG* expression. In both the terminal ileum and colon, the percentage of epithelial cells expressing *GCG* dropped in the inflamed mucosa and was already altered in tissues that did not appear inflamed by endoscopic examination compared to healthy controls. This is in line with the data from IL-10^-/-^ and *Tnf^ΔARE^* mice and suggests either the presence of low-grade inflammation or a persistent shift in epithelial subtype composition even in the absence of acute inflammation.

Notably, also the proportion of *GCG* expressing cells among EECs was reduced, indicating GLP-1- producing L cells to be particularly affected by inflammation (**Figure 1L,M**). Technical advances have overturned the one cell-one hormone dogma for EECs (44, 45), and thus, L cells co-express GLP-1 with different hormones depending on their location. Peptide YY (PYY) is one of these hormones and shows an expression pattern in the mammalian gastrointestinal tract similar to that of GLP-1 (46). Corroborating the hypothesis of L cell susceptibility to inflammation, the proportion of *PYY*-expressing EECs also decreased under inflammatory conditions in patients, in both the terminal ileum (**Figure 1N**) and colon (**Figure 1O**). Mirroring these results, also in the murine ileitis model (*Tnf^ΔARE^*) and colitis model (IL-10^-/-^) *Pyy* expression was reduced in inflammation-affected intestinal tissues (**Figure 1N,O**).

### 2.2 Inflammatory conditions impair epithelial differentiation into EECs

The intestinal epithelium constantly renews, and a tight balance between cell proliferation, differentiation, and apoptosis/ cell shedding is required to sustain homeostasis. Alterations in intestinal epithelial cell differentiation, subtype composition, and cell functions are part of the pathology of intestinal inflammation (15, 47).

Thus, we performed a transcriptional analysis of isolated intestinal epithelial cells from moderately inflamed IL-10-deficient mice and Wt littermates. Expression of transcription factors involved in epithelial differentiation and marker genes of epithelial subpopulations indicated reduced terminal differentiation of epithelial cells under inflammatory conditions (**Figure 2A,B**). This is consistent with crypt hyperplasia and expansion of the transit-amplifying zone, both histologic hallmarks of inflammation. EECs arise from the secretory linage, which also comprises antimicrobial peptide-producing Paneth cells (in the small intestine), Paneth-like cells (in the large intestine), mucin-producing goblet cells, serotonin-producing enterochromaffin cells, and tuft cells involved in chemosensing (**Figure 2B**). Secretory cells are particularly affected by inflammatory processes. For example, we found reduced numbers and functionality of Paneth cells in parallel to aberrations of the stem cell niche in CD patients and *Tnf^ΔARE^* mice (9, 39). Furthermore, goblet cell-loss is a common feature of *C. rodentium*-triggered colonic inflammation (42) as well as genetically-driven colitis observed in IL-10^-/-^ mice (48). Accordingly, we found an mRNA expression pattern indicating diminished presence of secretory precursor cells and reduced marker gene expression for all secretory cells except enterochromaffin cells (**Figure 2A,B**). Immunohistochemical staining and quantification of positive cells confirmed decreased numbers of Doublecortin-like kinase protein 1 (Dclk1)^+^ tuft cells but stable numbers of serotonin containing (5-HT^+^) enterochromaffin cell under inflammatory conditions in IL-10^-/-^ mice (**Figure 2C,D**), suggesting mechanisms specifically sustaining this cell population under inflammatory conditions.

**Figure 2:**
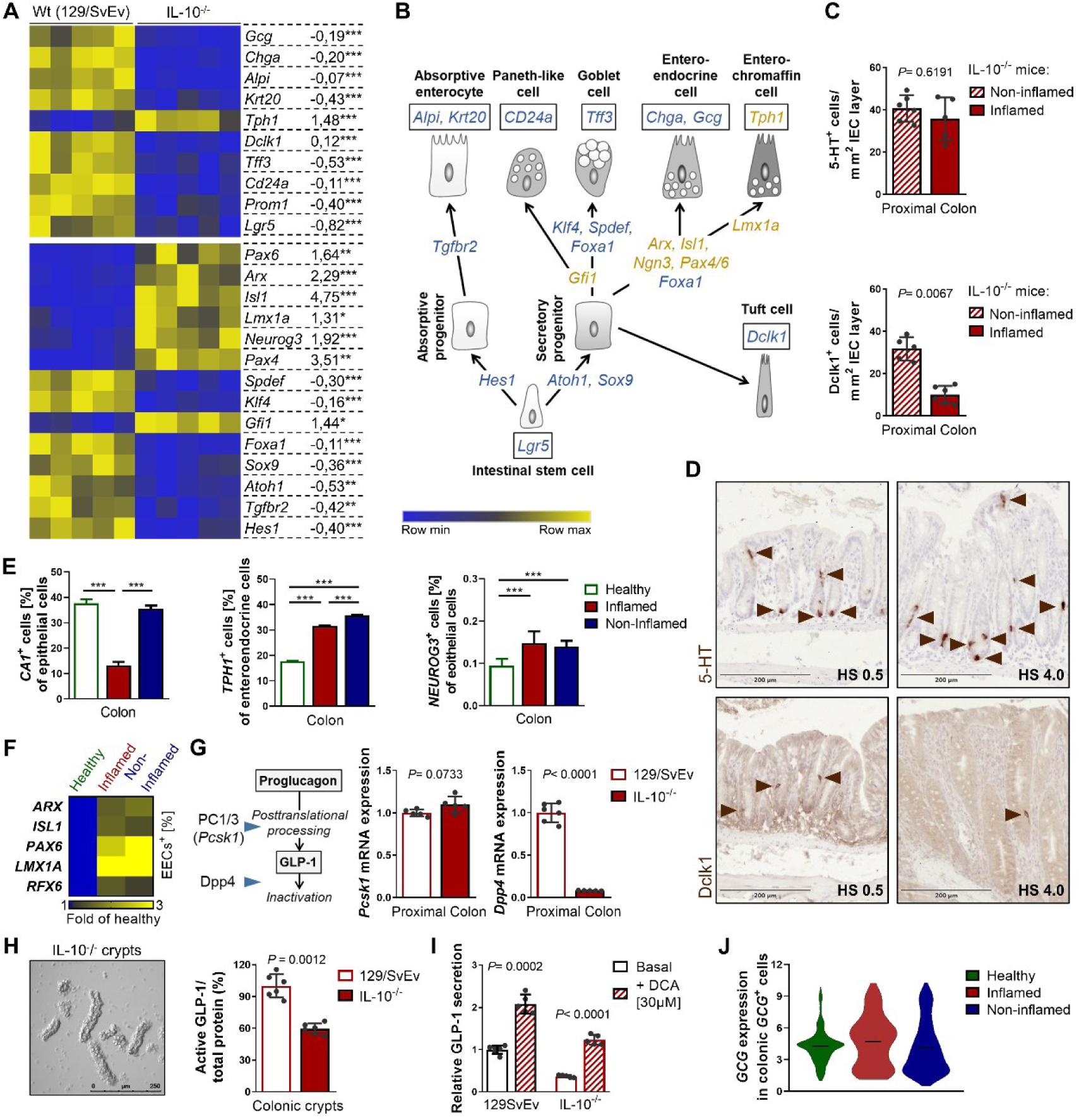
Inflammation selectively impairs epithelial differentiation but not L cell function. (**A**) Heatmap of a transcriptional profile of isolated intestinal epithelial cells from moderately inflamed (histological score (HS) 2-4) IL-10^-/-^ mice and wild-type (Wt) controls showing marker genes for epithelial subtypes and genes involved in epithelial differentiation. Fold changes in gene expression are given on the right side. Asterisks indicate significant differences \**P*<0.05, \*\**P*<0.01, \*\*\**P*<0.001. (**B**) Schematic representation of marker gene expression during differentiation and in epithelial subtypes. Blue = down-regulation, yellow = up-regulation under inflammatory conditions. (**C**) Quantification of 5-HT (serotonin)-positive enterochromaffin cells and Dclk1-positive tuft cells in colonic tissue sections of non-inflamed and inflamed IL-10^-/-^ mice. (**D**) Representative 5-HT and Dclk1 IHC stainings for the HS indicated. Arrows indicate cells positive for the respective marker. (**E**) Barplots showing the proportion of epithelial cells expressing *CA1*, a marker gene of mature colonocytes, (left) as well as *TPH1* expression within enteroendocrine cells (middle) in a scRNA- sequencing dataset of colonic intestinal epithelial cells from Crohn’s disease patients (35). Right: proportion of epithelial cells expressing the EEC progenitor marker *NEUROG3*. Bars represent mean+SD. (**F**) Heatmap showing the changes in the proportions of EECs expressing the indicated transcription factors involved in EEC differentiation. *P*<0.05 for all changes shown. One-way ANOVA followed by Tukey test, asterisks indicate significant differences \**P*<0.05, \*\**P*<0.01, \*\*\**P*<0.001. (**G**) qRT-PCR analysis for *Pcsk1* and *Dpp4*, enzymes regulating GLP-1 abundance (scheme, left), of ileal and colonic tissue from IL-10^-/-^ mice and wild-type (Wt) controls. (**H**) Representative picture of isolated colonic crypts from inflamed IL-10^-/-^ mice (left). Quantification of active GLP-1 (ELISA) in isolated colonic crypts from moderately inflamed IL-10^-/-^ mice and Wt controls (right). (**I**) Basal and deoxycholic acid (DCA)-induced secretion of active GLP-1 from primary colonic organoids derived from IL-10^-/-^ mice and Wt controls. Basal secretion from Wt colonic organoids is set as “1”. Bars represent mean+SD, n≥5 for all groups. Unpaired t tests, *P* values and are indicated. (**J**) Violin plot from the same dataset as in (**E**) showing the normalized expression height and distribution of *GCG* within *GCG*-expressing colonic epithelial cells. No significant differences are observed (one-way ANOVA).

Of note, a hierarchical cascade of transcription factors acts in a combinatorial fashion to steer differentiation of Neurogenin 3 (Neurog3)-expressing progenitor cells into the diverse EEC subtypes (46). Neurog3 itself is required to restrict secretory progenitors to the endocrine lineage (49) and was increased in intestinal epithelial cells from inflamed IL-10^-/-^ mice (**Figure 2A,B**). An early event in EEC differentiation is whether the cell commits to the enterochromaffin or peptidergic lineage. This decision is regulated by the transcription factor Regulatory factor 6 (Rfx6), which promotes expression of peptidergic hormones while repressing 5-HT synthesis *via* LIM homeobox transcription factor alpha (Lmx1a) (50). Accordingly, Rfx6 acts upstream of late acting transcription factors Aristaless-related homeobox (Arx), Paired box gene (Pax) 6 and Insulin gene enhancer protein (Isl) 1, regulating enterohormone gene transcription and as such, controlling terminal differentiation of enteroendocrine cell subtypes (46). Consequently, loss of Rfx6 in the epithelium causes accumulation of Neurog3- positive enteroendocrine progenitors and expansion of 5-HT-positive enterochromaffin cells, accompanied by impaired differentiation of peptide-producing EECs (50). Although this resembles the situation observed under inflammatory conditions, expression of *Arx*, *Pax6*, *Isl1* and *Lmx1a* was increased under inflammatory conditions in intestinal epithelial cells from IL-10^-/-^ mice (**Figure 2A,B**)., arguing against Rfx6 as key driver of skewed differentiation.

Employing the same single-cell RNA sequencing data set (35) as before, a similar pattern was observed in CD patients. There was a reduction in colonic epithelial cells expressing the mature colonocyte marker *CA1* (*Carbonic anhydrase 1*) under inflammation (**Figure 2E**). Concomitantly, the proportion of EECs expressing the enterochromaffin cell marker *TPH1* (*Tryptophan hydroxylase1*) and the proportion of epithelial cells expressing *NEUROG3* increased in CD patients independent of the inflammatory status (**Figure 2E**). Further reflecting the gene expression pattern of inflamed IL-10^-/-^ mice, the proportion of EECs expressing *ARX*, *PAX6*, *Isl1, LMX1A* and *RFX6* were increased in CD patients (**Figure 2F**). Interestingly, a tight bi-directional interaction of L cells and enterochromaffin cells has been described, with L cells expressing serotonin receptor 4 (51) and enterochromaffin cells expressing GLP-1 receptor (52). Moreover, endocrine cell differentiation is facilitated through paracrine GLP-1 and serotonin-mediated signaling involving Neurog3 (53).

### 2.3 Remaining colonic L cells are functional

The transcription factors *Pax6* and *Isl1* directly bind consensus sequences on the *Gcg* promoter, indicating a sustained expression of *Gcg* in the reduced number of GLP-1^+^ cells in the epithelium. Following translation, proglucagon encoded by *Gcg* is cleaved into active GLP-1 by prohormone convertase (PC) 1/3 in EECs. Subsequent to secretion, active GLP-1 is rapidly cleaved and inactivated by dipeptidyl peptidase-4 (DPP4) in target tissues, resulting in a half-life of only 2 minutes (54, 55) (**Figure 2G, scheme**). In accordance with the hypothesis that remaining EECs sustain GLP-1 expression, *Pcsk1* (encoding PC1/3) expression remained unaltered under inflammation in IL-10^-/-^ mice, whereas expression of *Dpp4* strongly decreased specifically at the site of inflammation (proximal colonic tissue) (**Figure 2G**), suggesting compensatory mechanisms to maintain GLP-1 signaling. Corroborating these results, we found reduced but substantial amounts of active GLP-1 in primary colonic crypts derived from IL-10^-/-^ mice (**Figure 2H**). Furthermore, L cells remained functional as primary colonic organoids derived from IL-10^-/-^ mice showed a secretory response towards stimulation with the bile acid deoxycholic acid (DCA) that did not significantly differ from that of Wt organoids (3.3 ± 0.36 *vs* 2.1 ± 0.28 fold increase), despite overall reduced secretion of GLP-1 per living cell (**Figure 2I**). Likewise indicating reduced but functional L cells, the average expression of *GCG* within colonic *GCG*^+^ cells remained unaltered in the single-cell RNA sequencing data set (35) in CD patients compared to healthy controls (**Figure 2J**).

### 2.4 Inflammatory cytokines and microbial structures affect *Gcg* expression

To investigate a potential contribution of pro-inflammatory cytokines and microbial structures to the inflammatory environment fostering L cell loss, we employed intestinal organoids derived from Wt mice. First, we tested whether microbial structures including bacterial muramyl dipeptide (MDP) and lipopolysaccharide (LPS) or the pro-inflammatory cytokines interferon γ (IFN) and TNF would induce GLP-1 secretion (**Figure 3A**). MDP is a substrate of PEPT1, known to mediate di/tripeptide-induced GLP-1 secretion in L cells and can be sensed intracellularly by nucleotide-binding oligomerization domain (NOD)-like receptors that belong to the innate immune system (56). SNPs in both *Pept1* and *Nod2* are known risk factors for IBD (57, 58). Yet, neither MDP nor treatment with a mixture of IFN/TNF led to a detectable secretion of GLP-1 despite responsiveness of organoids towards the bile acid deoxycholic acid (DCA) (**Figure 3B**). In contrast, LPS acted as a weak GLP-1 secretagogue, in accordance with literature proposing TLR4-dependent GLP-1 secretion upon LPS exposure by L cells as pathogen sensing mechanism during intestinal injury (59). A mixture of forskolin/3-isobutyl-1-methylxanthine (F/I) served as a control for maximal hormone output (**Figure 3B**).

**Figure 3:**
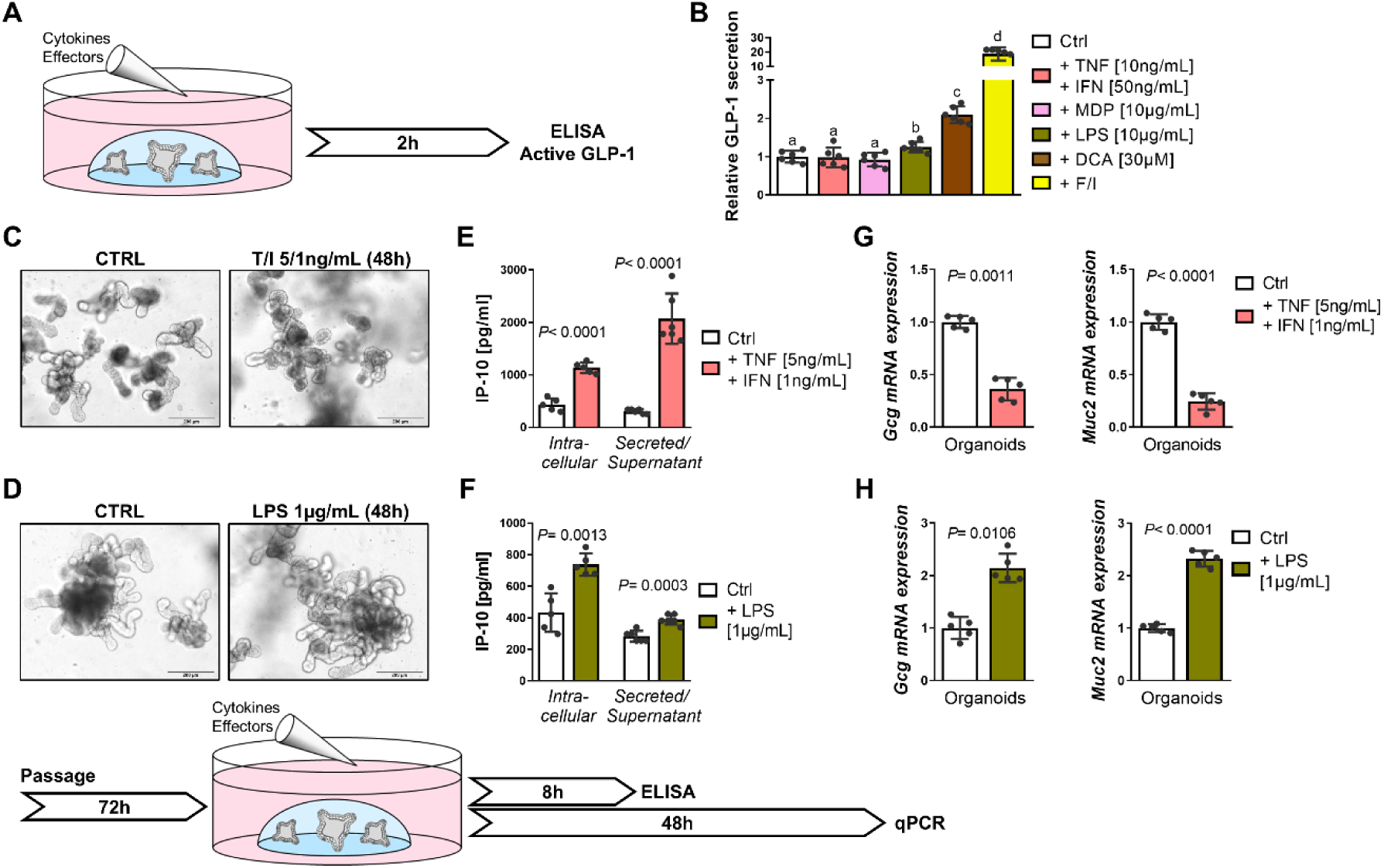
Inflammatory stimuli and microbial structures differentially affect *Gcg* gene expression and GLP-1 secretion. (**A**) Schematic representation of the experimental setup for (**B**), secretion of active GLP-1 (ELISA) from wild-type (Wt) intestinal organoids stimulated with bacterial-derived Lipopolysaccharide (LPS) or Muramyl dipeptide (MDP), or pro-inflammatory cytokines tumor necrosis factor (TNF) and Interferon gamma (IFN), respectively. Deoxycholic acid (DCA) serves as physiological positive control, forskolin/3-isobutyl-1-methylxanthine (F/I) induces maximal secretion. (**C,D**) Representative pictures of Wt intestinal organoids stimulated with TNF/IFN (**C**) or LPS (**D**) showing no growth impairment and schematic representation of the experimental setup. (**E,F**) Intracellular protein levels and secretion of the pro-inflammatory cytokine IP-10 in Wt intestinal organoids stimulated with (**E**) TNF/IFN or (**F**) LPS, respectively. (**G,H**) qRT-PCR analysis for *Gcg* and *Muc2* of intestinal organoids stimulated with (**G**) TNF/IFN or (**H**) LPS, respectively. (**B, E-H**) Bars represent mean+SD, n≥5 for all groups. Unpaired t tests except (**B**) one-way ANOVA followed by Tukey test. *P* values are indicated; a, b, c, d are significantly (*P* < 0.05) different from each other.

In a second approach, organoids were stimulated with low concentrations of TNF/IFN and LPS for 48h to determine the effect on epithelial differentiation. The chosen concentrations did not impair organoid survival or growth after 48h (**Figure 3C,D**), but robustly induced Interferon gamma-induced protein 10 (IP-10) in cell lysates and supernatants after 8h (**Figure 3E,F**), indicating reactivity towards stimulation. Treatment for 48h with TNF/IFN resulted in diminished expression of *Gcg* and *Mucin 2* (*Muc2*), a marker for goblet cells, recapitulating *in vivo* inflammatory conditions (**Figure 3G**). In contrast, LPS treatment increased *Gcg* and *Muc2* expression (**Figure 3H**), which supports the hypothesis that persistent activation of L cells stimulates L cell differentiation in a paracrine manner (60) and data showing LPS-mediated upregulation of mucin expression and secretion in goblet cells (61).

### 2.5 Inflammation-associated impairment of mitochondrial metabolism is associated with reduced enteroendocrine cell markers

A potential mechanism by which TNF/IFN and inflammation might affect L cell numbers is by affecting cellular metabolism and particularly mitochondrial function, including oxidative phosphorylation (OXPHOS) (15, 62, 63). Previously, we established a link between impaired mitochondrial function and altered epithelial cell differentiation in the context of intestinal inflammation (9, 14, 64, 65). In particular, we found an inflammation-related metabolic fingerprint in ileal epithelial cells comprising a shift from OXPHOS to glycolysis and overall reduced cellular ATP level in *Tnf^ΔARE^* mice. These metabolic changes were paralleled by induction of mitochondrial stress signaling pathways and aberrances of the intestinal stem cell niche, including loss of Paneth cells (9). On the basis of these results we screened genes indicative for different inflammation-relevant cellular metabolic pathways and metabolic genes known to affect the intestinal stem cell niche (16, 65) in isolated intestinal epithelial cells from moderately inflamed IL-10-deficient mice and Wt littermates (**Figure 4A**). Reflecting previous observations, transcriptional analysis indicated holistic metabolic changes in colitis, including reduced mitochondrial metabolism, enhanced glycolysis and activation of autophagy (66). Genes involved in the dynamic metabolic adaptions required for maintaining homeostasis of the ISC niche (including *Pdk4, Sgk1, Hmgcs2* and *Mtor*) were consistently down- regulated (**Figure 4A,B**). Concomitantly, cellular ATP levels were reduced and expression of *Trib3*, a marker for activation of the mitochondrial unfolded protein response (MT-UPR), a signaling pathway related to disturbances of mitochondrial function, was enhanced (**Figure 4C,D**). Highlighting the strong link between inflammation-related epithelial metabolism and *Gcg* expression, mRNA levels of *Hk2* and *Ddit4*, genes involved in glycolysis and suppressing mammalian target of rapamycin (mTOR) activity (67, 68), negatively correlated with *Gcg* expression, while a positive correlation was observed for *Pck1* and *Hmgcs2*, genes involved in gluconeogenesis and ketogenesis, respectively (**Figure 4E**). Importantly, these finding were reflected by the single-cell RNA sequencing data set derived from human mucosal biopsies. The frequency of colonic epithelial cells expressing marker genes for mitochondrial stress signaling (*TRIB3*, *ATF5*) and reduced cellular ATP levels (*PRKAA1* encoding the catalytic subunit of AMPK) was increased in tissues from inflamed CD patients (**Figure 4F**). Concomitantly, the proportion of colonic epithelial cells expressing metabolism-associated genes *HK2* and *DDIT4* was elevated under inflammatory conditions, while the proportion of cells expressing *PCK1* and *HMGCS2* decreased (**Figure 4G**). In line with previous results, most changes in cellular expressions were also visible in epithelial cells derived from biopsies of CD patients that were considered non-inflamed, again pointing toward cellular metabolic alterations preceding overt tissue pathology or persisting in the absence of the same. In summary, these data indicate that, consistent with ileal inflammation (9), colonic inflammatory processes are also associated with mitochondrial stress and are paralleled by reprogramming of intestinal cellular metabolism, dampening OXPHOS, and fostering glycolysis. This metabolic fingerprint accompanies the decline in GLP-1 positive cell numbers.

**Figure 4:**
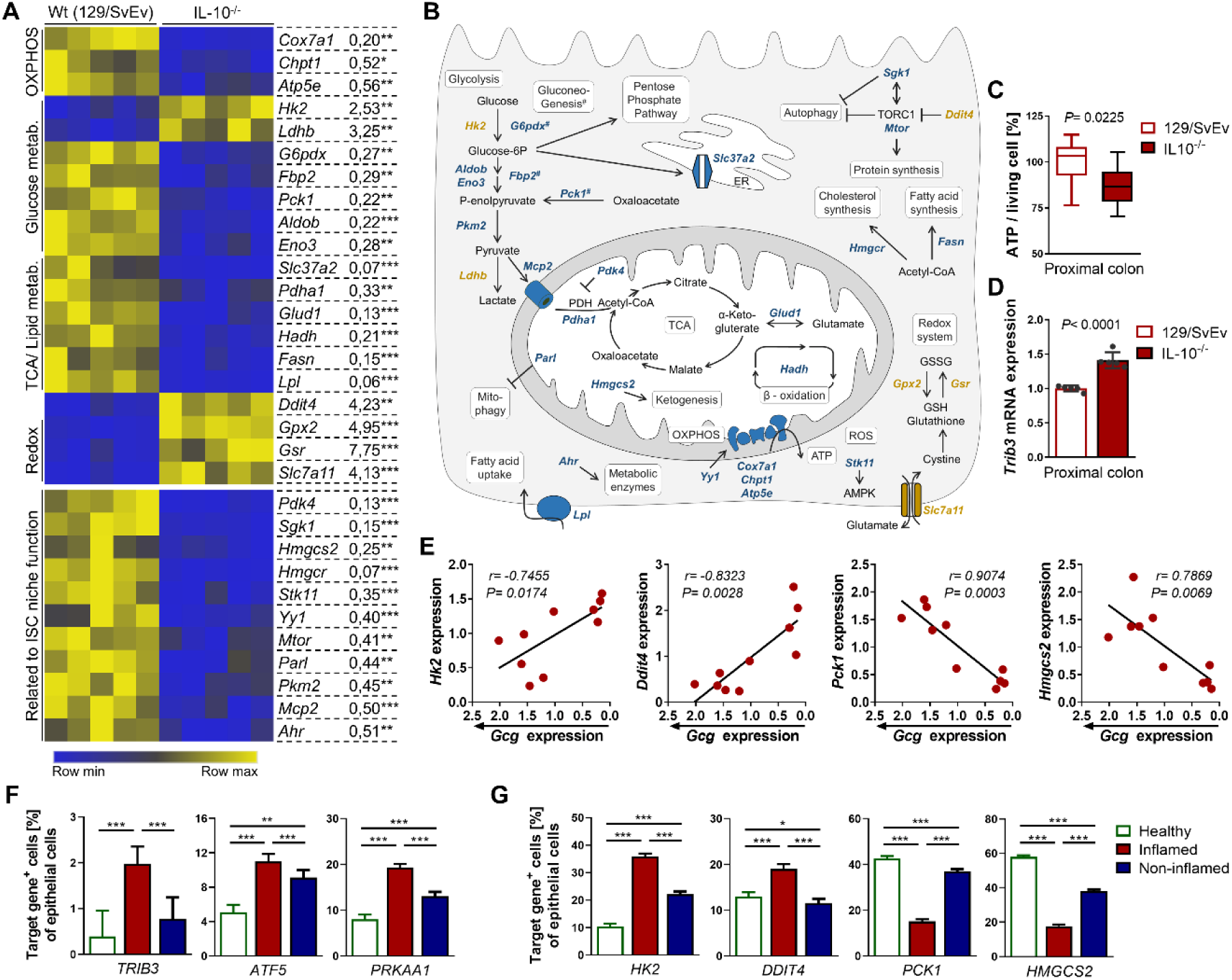
The inflammation-related metabolic profile is associated with GLP-1^+^ cell loss. (**A**) Heatmap of a transcriptional profile of isolated intestinal epithelial cells from moderately inflamed (HS 2-4) IL-10^-/-^ mice and wild-type (Wt) controls showing metabolism-associated genes involved in different metabolic processes and stem cell niche function. Fold changes in gene expression are given on the right side. Asterisks indicate significant differences \**P*<0.05, \*\**P*<0.01, \*\*\**P*<0.001. (**B**) Schematic representation of metabolic gene functions. Blue = down-regulation, yellow = up-regulation under inflammatory conditions. (**C**) ATP content of primary isolated colonic crypts from IL-10^-/-^ mice and Wt controls. (**D**) qRT-PCR analysis of isolated colonic epithelial cells for *Trib3*, a marker gene for mitochondrial stress signaling. (**E**) Correlation (Pearson) of metabolic gene expression with *Gcg* mRNA level in colonic epithelial cells isolated from IL-10^-/-^ mice. (**F, G**) Barplots showing the proportion of colonic epithelial cells expressing marker genes for mitochondrial stress signaling (*TRIB3*, *ATF5*) and AMP kinase activation (*PRKAA1*) or (**G**) target metabolic genes in a scRNA-sequencing dataset of colonic intestinal epithelial cells from Crohn’s disease patients (35) and SD of expression. One-way ANOVA followed by Tukey test, asterisks indicate significant differences \**P*<0.05, \*\**P*<0.01, \*\*\**P*<0.001. Bars represent mean+SD.

### 2.6 Mitochondrial impairment causes reduction of enteroendocrine cells

Intriguingly, deletion of the mitochondrial chaperone Hsp60 in murine colonocytes, triggering mitochondrial dysfunction and stress signaling, resembles the metabolic fingerprint of inflammation including up-regulation of *Hk2* and *Ddit4*, as well as down-regulation of *Pck1* and *Hmgcs2* (65). Elegantly demonstrating the causative contribution of metabolic disturbances to altered epithelial differentiation, this genetic model of primary mitochondrial dysfunction also shows a reduction in differentiated cells, including EECs, enterocytes, and tuft cells (but not enterochromaffin cells), as shown by single-cell RNA sequencing (65). To further establish mitochondrial impairment as a key driver of skewed epithelial differentiation processes, we used a mouse model with an epithelial-specific knockout of caseinolytic mitochondrial matrix peptidase (ClpP^ΔIEC^). ClpP contributes to mitochondrial proteostasis and knockout of ClpP triggers induction of Trib3 (69), Atf5 (70) and activation of AMPK (71). Tissue-specific ClpP deficiency in liver and skeletal muscle causes mitochondrial dysfunction including a decrease in mitochondrial respiratory supercomplexes (70). In contrast to Hsp60, whose non-inducible IEC-specific knockout (using *Villin1*Cre transgenic mice) is embryonic lethal (64), ClpP^ΔIEC^ mice are viable and exhibit normal growth and body weight development on a standard chow diet (data not shown), indicating less severe mitochondrial impairment. Epithelial loss of ClpP was confirmed by qPCR analysis of *ClpP* mRNA in isolated IECs (**Figure 5A**). Morphological features of the small intestine and colon including intestinal length, number of crypts (quantifications not shown), and crypt/villus length were similar to control *ClpP*^flox/flox^ littermates at the age of one year (**Figure 5B,C**). On the cellular level, proliferation (quantified by number of Ki67-positive cells per crypt) remained unchanged as well as the number of mucin-producing goblet cells (identified by PAS/AB staining) (**Figure 5C**). However, plasma concentration of citrulline, synthesized in the small intestinal epithelium as part of the urea cycle and serving as a functional readout of IEC mitochondrial function, was decreased in ClpP^ΔIEC^ mice, in line with previous results from Hsp60^ΔIEC^ mice (64) (**Figure 5D**). Additionally, Paneth cells from ClpP^ΔIEC^ mice contained fewer granules positive for the antimicrobial peptide lysozyme (Lyz^+^) (**Figure 5E,F**), despite overall Paneth cell abundance remaining unchanged (data not shown). Disturbances of AMP-packaging into granules and alterations in the granule exocytosis pathway manifest as disorganized and reduced numbers of cytoplasmic granules as well as a diffuse cytoplasmic lysozyme expression (8) (**Figure 5F, scheme**). Therefore, morphology of Lyz^+^ secretory granules is used as a functional marker for Paneth cell phenotype classification, and both inflammation and epithelial mitochondrial dysfunction have previously been shown to correlated with reduced Paneth cell granularity (8, 9). Importantly, numbers of GLP-1^+^ cells were diminished in the ileum and proximal colon in ClpP^ΔIEC^ mice (**Figure 5G**), indicating that mild mitochondrial dysfunction in the absence of tissue aberration or overt inflammation is sufficient to compromise Paneth cell function and reduce EEC differentiation.

**Figure 5:**
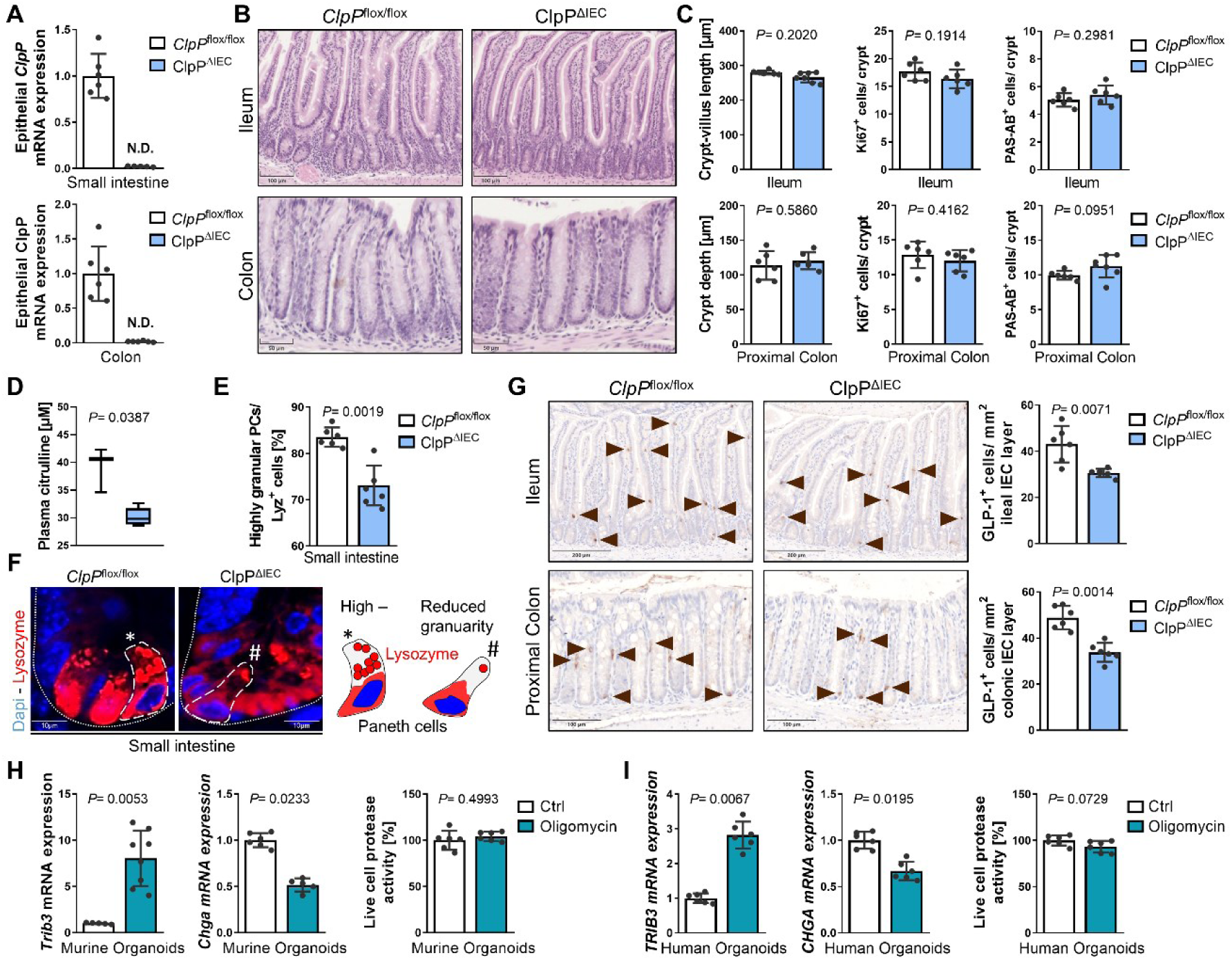
Mitochondrial dysfunction impairs epithelial differentiation into L cells. (**A**) qRT-PCR analysis for *ClpP* in small intestinal and colonic IECs from ClpP^ΔIEC^ mice and control *ClpP*^flox/flox^ littermates. (**B**) Representative pictures of ileal and colonic tissue sections, and (**C**) quantification of crypt/ villus length (left) actively proliferating cells marked by Ki67 (middle) and mucin-expressing goblet cells (right) in ileal and colonic tissue sections of ClpP^ΔIEC^ mice and controls. (**D**) Plasma citrulline level of ClpP^ΔIEC^ mice and controls. (**E**) Quantification of highly granular Lyz^+^ Paneth cells (PC) (≥2 granules) and (**F**) representative immunofluorescence staining of Lysozyme (red) counterstained with Dapi (nuclei, blue) in ClpP^ΔIEC^ mice and controls including scheme to visualize Paneth cell properties (right). (**G**) Quantification of GLP-1-positive cells and representative pictures of ileal and colonic tissue sections from ClpP^ΔIEC^ mice and controls immunohistochemically stained for GLP-1. Arrows indicate cells positive for GLP1. (**H**) qRT-PCR analysis of murine intestinal organoids treated with oligomycin (for 6h followed by 18h recovery) for *Trib3* and the enteroendocrine cell marker Chga. *Right*: measurement of living cells in organoid cultures after oligomycin treatment. (**I**) Same analysis as in (**H**) for human organoids. Bars represent mean+SD, n≥5 for all groups. Unpaired t tests, *P* values and are indicated.

To further validate that disturbances of mitochondrial metabolism skew epithelial differentiation, we used the OXPHOS inhibitor oligomycin *ex vivo* in intestinal organoids. Murine intestinal Wt organoids (**Figure 5H**) and human intestinal organoids (**Figure 5I**) were treated with sub-lethal doses of oligomycin for 6h and were subsequently allowed to recover for 18h. Blockade of OXPHOS induced mitochondrial stress signaling (MT-UPR) as evidenced by *Trib3*/ *TRIB3* upregulation and led to a reduction of the enteroendocrine marker gene *Chga*/ *CHGA*, in both murine and human organoids (**Figure 5H-I**). In summary, these data support the notion that impaired mitochondrial function in epithelial cells, as observed under inflammatory conditions, might be the driving force of reduced GLP- 1^+^ cell numbers.

## 3 Conclusion

While the mechanisms underlying hormone secretion from L cells and their functions in the context of digestion and metabolism are well studied, their role in the pathogenesis of intestinal inflammation is virtually unknown.

Quantifying GLP-1^+^ cells in four mouse models of ileitis and colitis and characterizing two independent cohorts of each *Tnf^ΔARE^* mice and IL-10^-/-^ mice, we found a consistent reduction in GLP-1 expressing cells confined to the site of active disease and correlating to the degree of inflammation. Importantly, we confirmed the reduced expression of the GLP-1-encoding gene *GCG* in a publicly available single-cell RNA sequencing data set of ileal and colonic mucosal biopsies derived from healthy controls as well as inflamed and non-inflamed tissues from CD patients (35). Hence, the loss of GLP-1^+^ cells appears to be a general feature of both small and large intestinal inflammation. One major goal of this work was to give a comprehensive overview of different mouse models of intestinal inflammation and therefore, most tissue samples used were generated in the context of previous studies. This explains the lack of GLP-1 measurements in plasma, a clear limitation that has to be overcome in subsequent studies. However, previous data do not indicate a reduced glucose tolerance in IL-10 deficient mice (66). This could be attributed to location-specific functions that have been proposed for GLP-1, with ileal GLP-1 stimulating insulin release in response to nutrients (72) and colonic GLP-1 exerting rather local effects, *i.e.*, slowing gastrointestinal transit time (73).

Regarding IBD patients, available data for plasma GLP-1 levels remain inconsistent (27). This may be due to methodological differences (74), but also biological challenges, including the heterogeneity of IBD pathologies and the fact that only about 10–15% of secreted GLP-1 reaches the systemic circulation in its active form (54, 55). Moreover, total changes in hormone secretion depend on the severity of inflammation and the extent of tissue affected and therefore, might be too minor to be recovered systemically. Beyond their classical endocrine function, being secreted into the bloodstream to act on receptor-expressing cells in distant target tissues, EEC-derived hormones also signal to neighboring cells in the epithelial layer and the lamina propria. Thus, even if systemic changes in GLP- 1 levels might not be readily detectable, altered paracrine signaling of GLP-1 could have profound effects on enterochromaffin cells (as mentioned above) (51–53), intraepithelial lymphocytes (29) and afferent neurons, in turn impacting intestinal motility, mucosal immune responses and feeding behavior under inflammatory conditions (75).

In particular, a subset of EECs termed “neuropod cells”, capable of forming synapses with neurons innervating the small intestine and colon, has gained increasing attention (75). Acting as sensory epithelial cell, neuropod cells form a neuroepithelial circuit that transduces signals between gut and brain (76). Terminal fibers of the vagus nerve, among others, express receptors for multiple EEC hormones and also enterochromaffin cell-derived serotonin acts on intestinal neurons (77). For instance, the effects of GLP-1 on food intake and glucose homeostasis are, at least in part, neurally mediated (78, 79), and co-release of GLP-1 and ATP from neuropod cells has been shown to elicit a vagal response (80). These neuroepithelial signals are thought to participate in sensing of nutritional value and the regulation of satiety, reward of food and peristaltic activity (75).

Furthermore, EECs are an integral part of the intestinal epithelial barrier, interacting with the microbiome as well as the mucosal immune system. Sensing the luminal and lamina propria environment, EECs not only react to bacterial structures like LPS through pattern recognition receptors (72, 81, 82), and bacterial metabolites such as SCFA and indole (83), but also express TNF receptor 1 (84). In addition to GLP-1 release, exposure to flagellin or lipopolysaccharide (85, 86) also results in expression of pro-inflammatory cytokines and evokes a gene response indicating EECs to participate in innate immune responses to commensals and pathogens (85). Highlighting the role of GLP-1 in regulating immune responses, gastrointestinal intraepithelial lymphocytes express the GLP-1 receptor (29), establishing a local enteroendocrine L cell-intraepithelial lymphocyte axis. GLP-1R signaling in intraepithelial lymphocytes selectively restrains local and systemic T cell-induced inflammation by suppression of effector functions, and modulates microbiota composition (87). These properties are of particular relevance in the context of IBD, where dysbiosis and dysregulated immune responses towards commensal constituents of the microbiota are thought to play a critical role in pathogenesis (14).

Hormone expression patterns of L cells differ considerably depending on their location in the intestine (88–90) and GLP-1 is co-expressed with other hormones including GLP-2, PYY, glucose-dependent insulinotropic polypeptide GIP, cholecystokinin (CCK) and secretin. Thus, the reduction of GLP-1 expressing cells under inflammatory conditions in the intestine should also profoundly affect signaling of additional gut hormones. Indeed, next to *GCG*, we also found a reduction in *PYY* expressing cells, which is in agreement with a study using colonic epithelial cells from treatment-naïve adult CD patients (47). With regard to mucosal homeostasis, GLP-2, posttranslationally processed from proglucagon like GLP-1 and secreted in equimolar amounts (91), is of particular interest. GLP-2 is an epithelial growth factor implicated in tissue healing following injury, improving barrier function and restoring epithelial homeostasis. Furthermore, GLP-2 possesses anti-inflammatory properties and sustains Paneth cell function (92). Consequently, it has been shown to ameliorate experimental colitis in several animal models (93–96) and a GLP-2 analog was proven effective in the treatment of active moderate to severe CD (97).

Importantly, we link the loss of GLP-1^+^ cells to an inflammation-related epithelial metabolic signature that was previously associated with impaired Paneth cell and goblet cell differentiation (9, 14) and that is reflected in colonic epithelial cells of CD patients. In accordance with the pivotal role of mitochondria in steering differentiation processes by balancing OXPHOS and glycolysis, compromised mitochondrial function in ClpP^ΔIEC^ mice, as well as *ex vivo* inhibition of OXPHOS in organoid cultures, was sufficient to impair EEC development.

This is in line with previous data (9, 98) and points toward compromised mitochondrial function as underlying cause of altered epithelial cell composition during the onset of inflammatory processes. Underlining the translatability of findings to the human disease condition, a downregulation of the ketogenesis pathway, as expected by a drop in *HMGCS2* expression, has been described for colonocytes of CD patients (35) and in line, reduced levels of the ketone body β-hydroxybutyrate were observed in the colonic mucosa of IBD patients which correlated with IBD activity index (99). Of note, a similar pattern of *HK2* expression to that we found in the single-cell RNA sequencing data set of CD patients (lowest in healthy controls and gradual increase from non-inflamed to inflamed tissue samples from IBD patients) has recently been reported in another cohort of IBD patients (100).

Remarkably, analysis of the single-cell RNA sequencing data set CD patients revealed metabolic changes and signs of mitochondrial stress signaling as well as changes in epithelial subtype composition even in mucosal tissues that were considered non-inflamed by endoscopy. Considering the chronic remittent pathology of CD with phases of remission and flares of inflammation, these cellular metabolic changes might render the mucosa susceptible for additional inflammatory triggers. Consequently, improving metabolic fitness and metabolic reprogramming have been identified as new therapeutic targets combating IBD by restoring functionality of IEC subtypes and enforcement of the epithelial regenerative capacity (14). Although so far rather neglected as player in intestinal inflammation, EECs might also profit from such metabolic interventions.

Recently, the influence of GLP-1 receptor agonists on the disease course of CD and UC patients with concomitant type 2 diabetes was investigated in a large-scale cohort study. The results suggest a potential therapeutic advantage of GLP-1 receptor agonists in managing IBD, by reducing hospitalizations, surgeries, and mortality (101). These beneficial outcomes underline the need for more research to scrutinize the role of GLP-1 in the pathogenesis of intestinal inflammation. However, given the numerous properties of GLP-1, it is likely that it affects inflammatory processes in the mucosa and disease-related symptoms on multiple levels (**Figure 6**). Thus, EECs and GLP-1-based therapies could represent an independent, promising strategy for intestinal inflammatory diseases.

**Figure 6:**
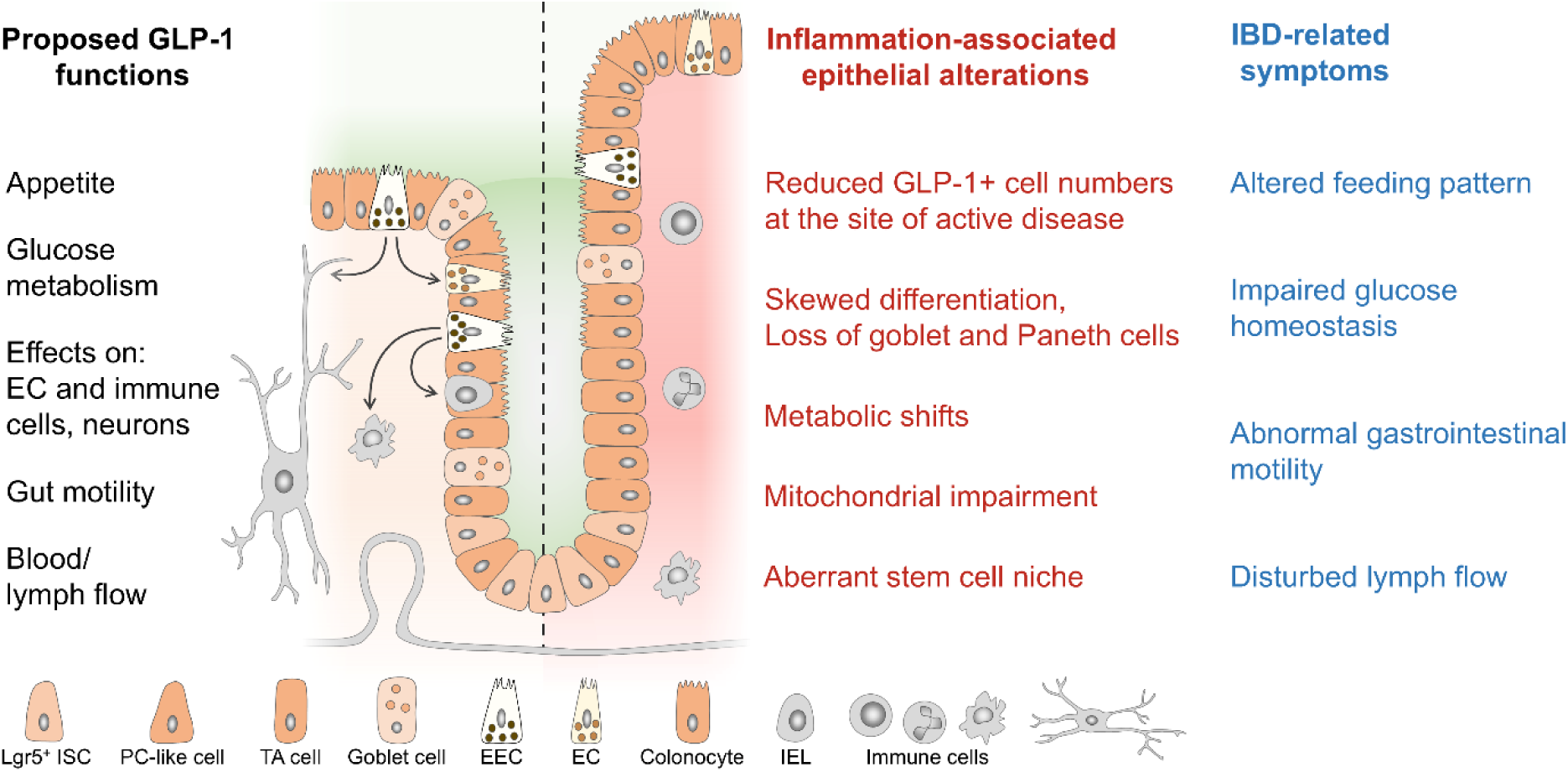
Schematic representation of GLP-1 functions, inflammation-associated epithelial alterations and IBD symptoms. Schematic representation of GLP-1 functions, inflammation-associated epithelial alterations and IBD symptoms.

## 5 Methods

All authors had access to the study data and had reviewed and approved the final manuscript.

### 5.1 Ethics statement

The maintenance and breeding of mouse lines and all experiments were approved by the Institutional Animal Care and Use Committee (IACUC), University of North Carolina at Chapel Hill, NC or approved by the Committee on Animal Health and Care of the local government body of the state of Upper Bavaria (Regierung von Oberbayern; approval numbers 55.2-1-54-2531-164-09, 55.2-1-54- 2531-99-13, 55.2-1-54-2532-217-2014) and performed in strict compliance with the EEC recommendations for the care and use of Lab. Anim. (European Communities Council Directive of 24 November 1986 (86/609/EEC)). For generation of human organoids, the use of surgically resected human tissue samples was approved by the Ethics Committee of the Medical Faculty of TUM and all participants in this study gave their written informed consent.

### 5.2 Animals

In accordance with 3R principles, most samples analyzed in this study for IHC-based quantification of intestinal cells were obtained from our tissue biobank and generated in the context of previously published studies (9, 36–38, 102).

**IL-10-deficient (IL-10^−/−^) mice** on 129Sv/Ev background or C57BL/6 background, ***Tnf^ΔARE^* mice** and **ClpP^ΔIEC^ mice** were housed (12h light/ dark cycles, 24-26°C) under specific pathogen-free (SPF) conditions according to the criteria of the Federation for Laboratory Animal Science Associations (FELASA) and bred for several generations in the animal facility. *Tnf^ΔARE^* mice were initially provided from Case Western University, Cleveland. *ClpP*^flox/flox^ mice were generated by Taconic-Artemis (Cologne, Germany) in close consultation with our lab, similar to described previously by others(103). To generate epithelial cell-specific ClpP knockout (ClpP^ΔIEC^) mice, *ClpP*^flox/flox^ mice were crossed to Cre transgenic *Villin1*Cre mice (C57Bl/6N), originally provided by Klaus-Peter Janssen (Klinikum Rechts der Isar, TU München). Mice received a standard chow diet (autoclaved V1124-300; Ssniff, Soest, Germany) and autoclaved water ad libitum. IL-10^−/−^ mice, *Tnf^ΔARE^* mice and Wt littermates were sacrificed at 16-18 weeks of age (9, 36, 37). Male ClpP^ΔIEC^ mice were analyzed at one year of age. Animals were killed by CO_2_ inhalation.

**Adoptive T -cell transfer** experiments were performed as previously described (38, 102). Briefly, lymphocyte-deficient SPF 129 SvEv recombination-activating gene (Rag) 2^-/-^ mice were reconstituted at 8 weeks of age by intraperitoneal injection of 5 x 10^5^ CD4^+^ T cells. CD4^+^ CD25^-^ and CD4^+^ CD25^+^ T cells for transfer were obtained from splenocyte suspensions of SPF Wild type mice using a magnetic bead-based CD4^+^ T cell-isolation kit or the CD4^+^ CD25^+^-Regulatory T cell-isolation Kit (Miltenyi Biotec) according to the manufacturer’s instructions. 4 weeks later, mice were killed by CO_2_ anesthesia followed by cervical dislocation. Reconstitution of mice was confirmed by flow cytometric analysis. Non-reconstituted Rag2^−/−^ mice served as controls.

#### Citrobacter rodentium infection

The *C. rodentium* strain DBS100 (ATCC 51459; American Type Culture Collection) was used for infection as previously described (104). *C. rodentium* was grown overnight at 37°C in Luria broth. C57BL/6J mice were infected by oral gavage with 0.5 mL of phosphate-buffered saline (PBS) containing approximately 1×10^9^ CFU of C. rodentium. To assess the clearance of C. rodentium, fecal pellets were collected from individual mice, homogenized in PBS, serially diluted and plated onto selective MacConkey agar. After an overnight incubation at 37°C, colonies were counted based on the size and distinctive appearance of *C. rodentium* colonies.

### 5.3 Mouse tissue processing and histopathological analysis

The intestine was removed immediately after euthanization, trimmed free of adjacent tissue and cleaned of stool. Ileal or proximal colonic intestinal tissue was either immediately fixed for preparation of cross sections or cut open and processed as ‘swiss role’. Tissues were fixed in 4% phosphate-buffered formalin for 48h, dehydrated and embedded in paraffin (FFPE). For histopathological assessment, 5 μm sections were stained with hematoxylin (of Mayer) and 0.2% eosin (ethanolic solution; both Medite) in an automated staining machine (Leica). Images were acquired by a PreciPoint M8 microscope (Precipoint). Ileal and colonic sections were blindly assessed for mononuclear cell infiltration into lamina propria, crypt hyperplasia, goblet cell depletion and ulcer formation resulting in a score from 0 (not inflamed) to 12 (highly inflamed) as described previously (48).

### 5.4 Morphometric measurements of intestinal tissues

The crypt depth and number of crypts per area were analyzed in H&E stainings of intestinal tissue sections. In fifty well-oriented crypts, the length from the crypt base to the crypt top or villus tip, respectively, was measured using the ViewPoint Light software, and the average length was calculated for each section. To determine the number of crypts per 100µm, the number of crypts within 100µm intestinal tissue length was counted in well oriented tissue areas using the ViewPoint Light software.

### 5.5 Alcian blue / Periodicacid-Schiff (PAS-AB) staining and quantification of goblet cells

FFPE tissue sections were deparaffinized, rehydrated, and stained with alcian blue (Fisher, Dreieich, Germany) for detection of acidic mucins (0.5% v/v in 3% acetic acid, pH=2.5, 5 min) in goblet cells. Sections were then treated with periodic acid (0.5% v/v, 10 min) and co-stained with Schiff’s reagent (Sigma- Aldrich, Taufkirchen, Germany) for neutral mucins (15 min). Nuclei were counterstained with hematoxylin, and tissue sections were dehydrated and mounted with DPX new (Merck KGaA, Darmstadt, Germany). Slides were scanned and further analyzed using a PreciPoint M8 microscope (Precipoint, Freising, Germany). The number of goblet cells was counted based on Alcian blue/Periodicacid-Schiff (PAS-AB) staining. Fifty well-oriented crypts were selected per mouse, and PAS^+^ cells were counted. The number of PAS^+^ cells was calculated per crypt/ per mouse.

### 5.6 Immunohistochemistry

Immunohistochemical (IHC) stainings were performed on 5 μm FFPE tissue sections as described previously (64). Briefly, FFPE tissue sections were deparaffinized and rehydrated in an automated staining machine (Leica). After heat-mediated antigen retrieval (boiling in 10mM citrate buffer (pH=6) in a microwave (23min; 900 watt)), sections were incubated for 10min with 3% H_2_0_2_ (Sigma-Aldrich). Subsequently, sections were blocked, and a primary antibody was applied overnight at 4°C. After several washing steps, sections were incubated with the secondary antibody (horseradish peroxidase coupled; 1h, room temperature). Antigen detection was performed using 3,3′-diaminobenzidine (DAB) or enhanced DAB (Sigma Aldrich), and nuclei were counterstained with hematoxylin. Slides were mounted with xylol-based mounting medium (Roti®-Histokitt). Stainings were scanned and using a PreciPoint M8 microscope (Precipoint). Antibodies and dilutions are given in **Table 1 and 2**.

**Table 1:**
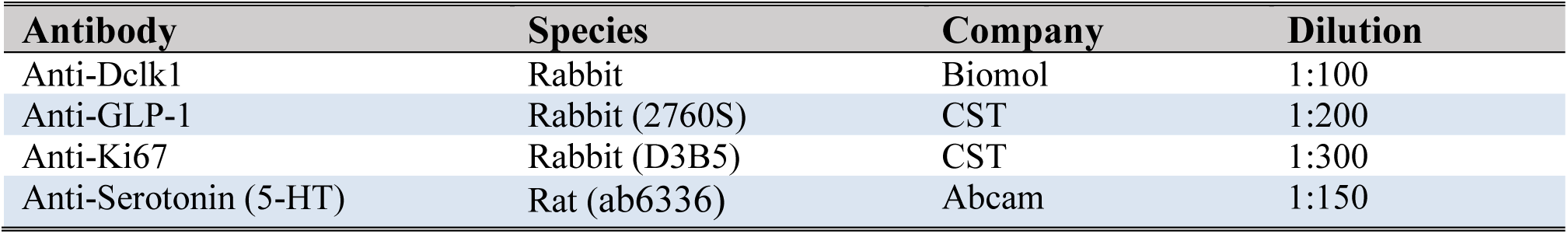
Primary antibodies used for IHC.

**Table 2:** Secondary antibodies used for IHC.

| Antibody | Company | Dilution |
| --- | --- | --- |
| HRP conjugated donkey anti rabbit IgG | Dianova (711-035-152) | 1:300 |
| HRP conjugated goat anti rat IgG | Dianova (112-035-003) | 1:300 |

### 5.7 Immunofluorescence stainings and Paneth cell phenotype quantification

Immunofluorescence stainings for lysozyme were performed on 4 μm FFPE tissue sections as described previously(9). Antibodies used were: Anti-Lysozyme Rabbit Dako, Agilent, Santa Clara (1:1000), Alexa Fluor donkey anti rabbit 546 Life Technologies, Carlsbad, CA (1:200) and Dapi to counterstain nuclei, Sigma- Aldrich, Taufkirchen, Germany (1:1000). Stainings were visualized using the Flouview FV10i microscope (Olympus, Shinjuku, Japan). Lysozyme positive cells numbers were determined in ileal tissue sections. To assess the granularity of Lyz positive cells, all PCs contained in 50 well oriented crypts were analyzed. Lyz positive cells were considered as highly granular if ≥2 Lyz positive granules were visible within the cell. The quantification was performed in a blinded manner.

### 5.8 Quantification of cell numbers

At least three non-consecutive sections from each mouse were used for quantification of GLP-1, Dclk1, Ki67 or 5-HT positive cells. Well-oriented (visible open crypts/ crypt-villus axis) tissue areas were selected and epithelial cells positive for the respective marker were counted. The number of positive cells was calculated over the total epithelial area quantified using the ViewPoint Light software (Precipoint). For individual mice, the number of positive cells was calculated as mean of quantified sections and areas. For quantification of Ki67 expressing cells, cells positive for Ki67 were counted in 50 well-oriented crypts per mouse and the average number of Ki67 positive cells was calculated per crypt/ per mouse. The quantification was performed in a blinded manner.

### 5.9 Isolation of primary IECs

Primary IECs were purified as previously described (64). Approximately 6 cm of intestine were inverted on a needle, vortexed vigorously and incubated (37 °C, 15 min) in DMEM containing 10% fetal calf serum (Biochrom), 1.0% Glutamine, 0.8% antibiotics/antimycotics (all Sigma-Aldrich) supplemented with 1 mM dithiothreitol (Roth). The IEC suspension was centrifuged (7 min, 300*g*, RT) and the cell pellet was re-suspended in DMEM containing fetal calf serum, l-glutamine and antibiotics. The remaining tissue was incubated in 20 ml PBS (10 min, 37 °C) containing 1.5 mM EDTA (Roth). After vortexing, the tissue was discarded and the cell suspension from this step was centrifuged as mentioned above. Cells obtained in both steps were combined and primary IECs were purified by centrifugation through a 20%/40% discontinuous Percoll gradient (GE Healthcare) at 600*g* for 30 min. IECs were immediately snap frozen.

### 5.10 Primary crypt isolation and intestinal organoid culture

Primary intestinal crypts from mouse (ileum, colon) and human (from macroscopically healthy tissue of surgical specimen of the ileum) were isolated by tissue incubation in 2mM EDTA (Fisher) or Gentle Cell Dissociation Reagent (STEMCELL Technologies), respectively, and cultured as described elsewhere (64, 105, 106). For organoid culture, murine crypts were embedded in 25µl matrigel (BD Biosciences) and cultured in 48 well plates. Murine organoids were grown in crypt culture medium (CCM), advanced DMEM/F12 medium (Gibbco) containing 2 mM GlutaMax (Gibbco), 10 mM HEPES, penicillin, streptomycin and amphotericin (all Sigma- Aldrich) supplemented with N2, B27 (both Gibbco), 1 mM N-acetylcystein (Sigma- Aldrich), 50 µg/mL EGF (ImmunoTools), 100 µg/mL noggin and 0.5 µg/mL R-spondin 1 (both PeproTec) or, when derived from colonic crypts, in Murine Intesticult medium (STEMCELL Technologies). For human organoid culture, 500 crypts were embedded in 30µl matrigel (BD Biosciences) per well, resulting in approx. 100 mature organoids, and cultured in 48 well plates. Human organoids were cultivated in Human Intesticult medium (STEMCELL Technologies) containing Wnt factors.

### 5.11 Intestinal organoid stimulation

For assessing the GLP-1 secretory capacity of IL-10^-/-^ enteroendocrine cells, primary colonic crypts were embedded in matrigel and cultured for 48h prior to stimulation to retain the inflammatory phenotype. For other experiments, murine and human organoids were used 4 to 6 days after passaging. Two days prior to the start of experiments, medium of human organoids was changed to CCM to reduce stem niche factors (106). When indicated, organoids were stimulated with TNF (5ng/mL) and IFN (1 ng/mL) or LPS (1µg/mL) for 48h, or treated with 2.5µM oligomycin (Sigma- Aldrich) for 6h and subsequently cultured in CCM for 18h, for gene expression analysis to assess differentiation. To measure IP-10 production and secretion, organoids were stimulated with TNF/ IFN or LPS in the same concentrations for 24h. To measure GLP-1 secretion, organoids were washed four times with HEPES- saline buffer (pH 7.4) containing BSA and DPP4 inhibitor. Subsequently, organoids were incubated for 2h in HEPES-saline buffer supplemented with either LPS (10µg/mL), MDP (10µg/mL), a combination of TNF (10ng/mL) and IFN (50 ng/mL), DCA (30 μM) (as physiological positive control), or a mixture of 10 μM Forskolin and 10 μM 3-isobutyl-1-methylxanthine (F/I, for maximal hormone output).

### 5.12 GLP-1 measurements

Organoid culture: after stimulation, the culture plate was placed on ice, supernatants were collected and immediately cold PBS was added to the organoid cultures. Supernatant samples were centrifuged (400 g) for 5 min at 4 °C and the supernatant was snap-frozen in liquid nitrogen. Organoids including Matrigel were resuspended in cold PBS and washed twice by centrifugation (600 g) for 10 min at 4 °C. The organoid pellet was dissolved in lysis buffer containing 150 mM NaCl, 50 mM Tris-HCl, 1% Igepal CA-630, 0.5% DCA and a protease inhibitor mix, for 20 min at 37 °C, and centrifuged (600 g) for 10 min at 4 °C. The supernatant was collected and snap-frozen. Primary crypts were lysed and processed accordingly. All samples were kept at −80 °C until analysis of incretin hormones. GLP-1 concentrations in supernatant and lysate were quantified using a Glucagon-Like Peptide 1 (Active) ELISA kit (Millipore). The amount of GLP-1 secreted by the cultures was calculated as a percentage of the GLP-1 content (secreted plus intracellular) and normalized to basal secretion. The content of active GLP-1 in primary crypts was normalized to total protein content (determined by Bradford assay) of samples and calculated as percentage of Wt (129/SvEv) colonic crypts.

### 5.13 IP-10 measurements

After stimulation, supernatants and organoids were collected according to the procedures described for GLP-1 measurement. The organoids were dissolved using a non-denaturating lysis buffer containing PMSF and a protease inhibitor mix. IP-10 concentrations were determined using an IP-10 ELISA kit (R&D Systems) according to the manufacturer’s instructions.

### 5.14 Measurement of living cells and cellular ATP content

Life- and dead- cell protease activity was measured using the MultiTox-Fluor Cytotoxicity Assay and ATP content was determined using the CellTiter-Glo Luminescent Cell Viability Assay (both from Promega), according to the manufacturer’s instructions. Fluorescence and luminescence were measured using the Tecan infinite M200 (Tecan Group Ltd.) and the i-control™ Microplate Reader software (Tecan).

### 5.15 Measurement of plasma citrulline

EDTA blood was taken during sampling from venacava. Quantitation of amino acids in EDTA plasma was performed using LC–MS/MS with aTRAQ labelling (aTRAQ reagent kit 4442671, AB SCIEX, Framingham, MA, USA). Quantitation was done according to the manufacturer’s instructions using 40 μl of serum sample. The analysis was performed on a triple quadrupole QTRAP3200 LC–MS/MS system (AB SCIEX) coupled to an Agilent 1260 Infinity Quaternary LC Pump (Agilent, Santa Clara, CA, USA).

### 5.16 RNA isolation, reverse transcription and quantitative real-time PCR

For mRNA isolation from primary IEC or organoids, the NucleoSpin RNAII kit (Macherey-Nagel) was used according to the manufacturer’s instructions. In the case of organoid culture, matrigel containing organoids was directly dissolved in RA1 buffer. Reverse transcription was performed as previously described (107). Quantitative Real-time PCR (qRT–PCR) was performed using the Light Cycler 480 system (Roche Diagnostics) and applying the Universal Probe Library system according to the manufacturer’s instructions. Primer sequences and probes are given in **Table 3**. Relative induction of gene mRNA expression was calculated based on Ct values using the expression of house-keeping genes *HPRT*/*Hprt* for normalization. Data are expressed as fold change over controls.

**Table 3:**
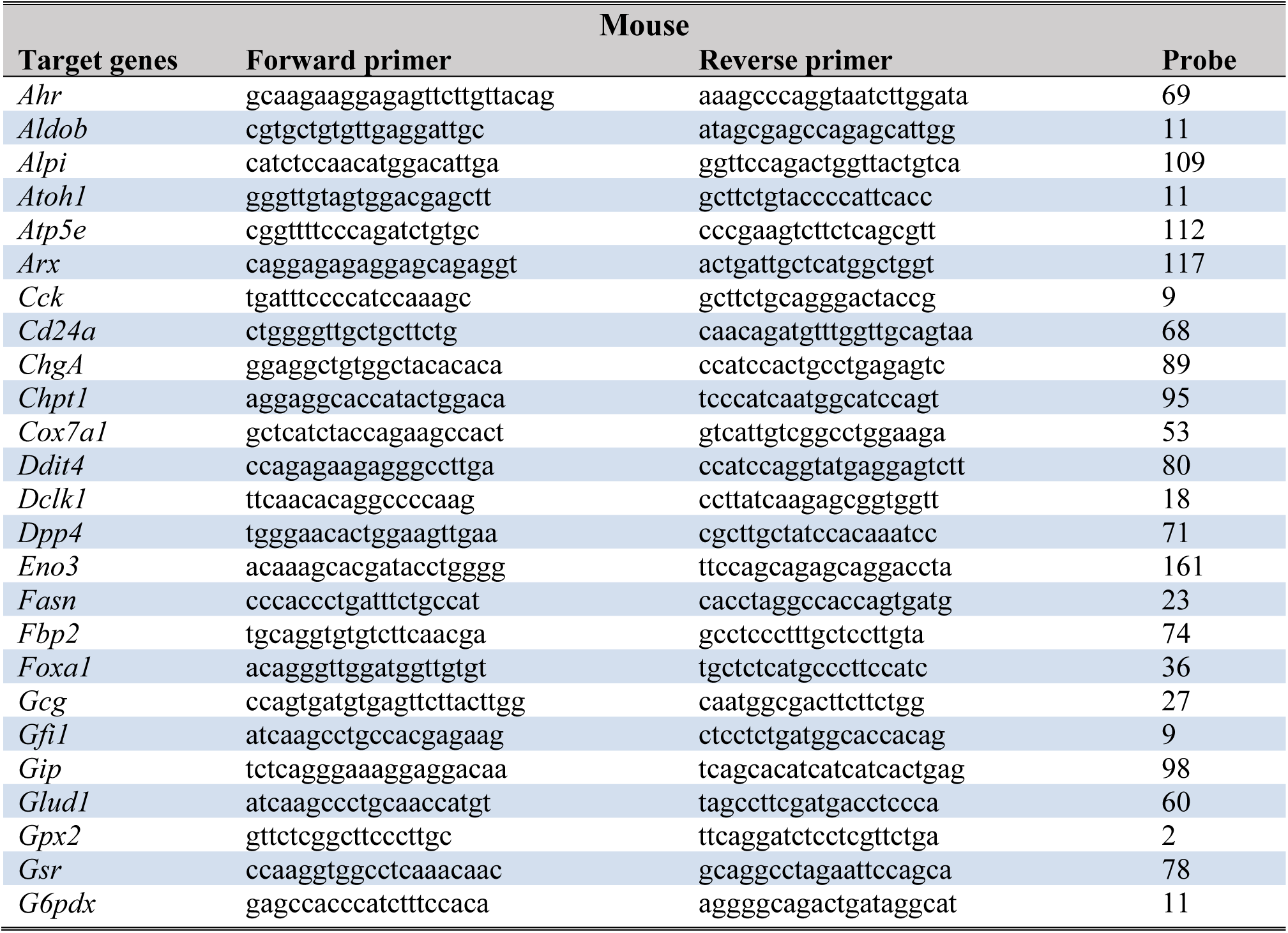
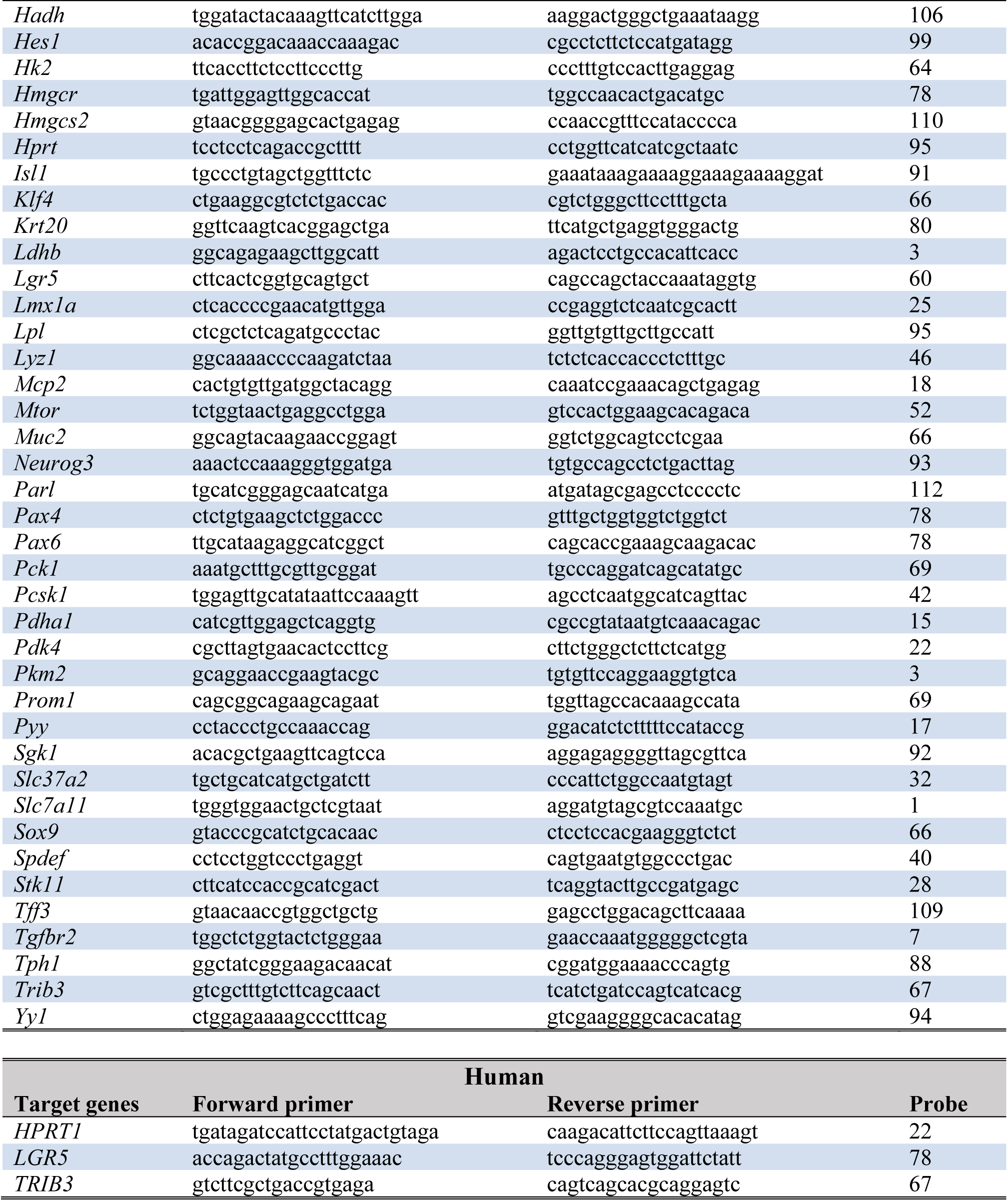
Primer sequences and probes.

### 5.17 Single-cell RNA sequencing and data visualization

The single-cell RNA sequencing dataset comprising intestinal epithelial cells from the terminal ileum and colon of healthy donors as well as inflamed and non-inflamed tissues from Crohn’s disease patients was obtained from the Human Cell Atlas portal (https://explore.data.humancellatlas.org/projects/cae461de-ecbd-482f-a5d4-11d607fc12ba) and initially published by Kong et al. (35). This data set comprises 720,633 cells across 71 donors with various inflammation statuses. We used pre-processed Seurat objects provided by the original authors, which included normalization, dimensionality reduction, and clustering. Analysis was performed in R (v4.3.3) using Seurat (v5.1.0). Cell type annotations and UMAP embeddings were directly extracted from the metadata. Ensembl gene IDs were converted to gene symbols using biomaRt (v2.58.2). Subsets of enteroendocrine cells were generated based on cell type annotations, and UMAP plots were used to visualize marker gene expression (e.g., *GCG*). For each subset, we calculated the proportion of cells expressing target genes (e.g., mitochondrial stress markers, metabolic genes) and computed average expression levels by inflammation status (healthy, inflamed, and non-inflamed), including standard deviation, sample size, and standard error of the mean. The complete analysis code is available at: https://github.com/Kate0107/scRNAseq-analysis-of-GLP-1-secreting-enteroendocrine-cells-in-the-context-of-intestinal-inflammation/blob/main/GLP1.R.

### 5.18 Statistical analysis and heatmap generation

Heatmaps were generated using Morpheus (https://software.broadinstitute.org/morpheus) or GraphPad Prism (GraphPad). Statistical analysis was performed using GraphPad Prism. Statistical tests used were: unpaired t-test for two group-comparisons or chi-square test or One-Way analysis of variances (ANOVA) followed by an appropriate multiple comparison procedure for comparisons comprising more than two groups. Correlation analysis was performed according to Pearson test. Data are expressed as mean ± SD, n ≥ 5 for all groups. Differences between groups were considered significant if P-values were < 0.05.

## Disclosures

The authors declare that the research was conducted in the absence of any commercial or financial relationships that could be construed as a potential conflict of interest.

## Preprint Server

https://www.biorxiv.org/content/10.1101/2025.03.05.641577v1

## Grant support

NIH P40OD01995, P30DK034987, P01DK94779 (RBS), Crohn’s & Colitis Foundation (RBS)

## Abbreviations

AMPK: 5’-prime-AMP-activated protein kinase
Atf/ ATF: Activating transcription factor
ATP: Adenosine triphosphate
Cck/ CCK: Cholecystokinin
CD: Crohn’s disease
ClpP: Caseinolytic mitochondrial matrix peptidase
Dclk: Doublecortin-like kinase protein
EEC: Enteroendocrine cell
Gcg/ GCG: Glucagon
GLP: Glucose like protein
Hmgcs2/ HMGCS2: 3-Hydroxy-3-methylglutaryl-CoA synthase
HS: Histoscore
IBD: Inflammatory bowel diseases
IEC: Intestinal epithelial cell
IL: Interleukin
Isl: Insulin gene enhancer protein
Mtor: Mechanistic target of Rapamycin
Ngn/NEUROG: Neurogenin
Pax: Paired box
PRKAA: Protein kinase AMP-activated catalytic subunit alpha
Pyy/PYY: Peptide YY
Rfx6/ RFX6: Regulatory Factor X 6
Tnf: Tumor necrosis factor
TPH: Tryptophan hydroxylase
Trib/TRIB: Tribbles pseudokinase
UC: Ulcerative colitis
Wt: Wild type

## Notes

### Competing Interest Statement

The authors have declared no competing interest.

### Summary of Updates

Changes in Figure 1, 2, and 5

